# Expansion of the *HSFY* gene family in pig lineages

**DOI:** 10.1101/012906

**Authors:** Benjamin M Skinner, Kim Lachani, Carole A Sargent, Fengtang Yang, Peter Ellis, Toby Hunt, Beiyuan Fu, Sandra Louzada, Carol Churcher, Chris Tyler-Smith, Nabeel A Affara

## Abstract

Amplified gene families on sex chromosomes can harbour genes with important biological functions, especially relating to fertility. The *HSFY* family has amplified on the Y chromosome of the domestic pig (*Sus scrofa*), in an apparently independent event to an *HSFY* expansion on the Y chromosome of cattle (*Bos taurus*). Although the biological functions of *HSFY* genes are poorly understood, they appear to be involved in gametogenesis in a number of mammalian species, and, in cattle, *HSFY* gene copy number correlates with levels of fertility.

We have investigated the *HSFY* family in domestic pigs, and other suid species including warthogs, bushpigs, babirusas and peccaries. The domestic pig contains at least two amplified variants of *HSFY*, distinguished predominantly by presence or absence of a SINE within the intron. Both these variants are expressed in testis, and both are present in approximately 50 copies each in a single cluster on the short arm of the Y. The longer form has multiple nonsense mutations rendering it likely non-functional, but many of the shorter forms still have coding potential. Other suid species also have these two variants of *HSFY*, and estimates of copy number suggest the *HSFY* family may have amplified independently twice during suid evolution. Given the association of *HSFY* gene copy number with fertility in cattle, *HSFY* is likely to play an important role in spermatogenesis in pigs also.

## Introduction

Sex chromosomes, and Y chromosomes in particular, are sites of frequent evolutionary innovation, due in part to the smaller population size of these chromosomes compared to autosomes, the lack of recombination on the Y, frequent and dramatic remodelling of the Y chromosome, and the accumulation of ampliconic sequences. During our collaborative project sequencing the pig X and Y chromosomes (Skinner et al, in submission), we became interested in a sequence that appeared repeatedly in the data being produced: the pig *HSFY*. The structure and organisation of the pig X and Y chromosomes are described in an attendant manuscript (Skinner et al. in submission), and in previous papers (Skinner et al. 2013; Groenen et al. 2012). Briefly, the Y chromosome long arm is highly repetitive, with all known single copy genes on the short arm. A central band of repetitive material is also found on the short arm at cytogenetic band Yp1.2.

Little is known about the biological function of *HSFY*. In humans, two functional *HSFY* copies are found in the AZFb region (Tessari et al. 2004; Shinka et al. 2004). Deletions in the AZFb region of the human Y are usually linked to problems with fertility in patients; however, microdeletions in the AZFb region affecting only *HSFY* do not seem to impair fertility (Kichine et al. 2012). The gene encodes a heat shock transcription factor, but it appears not to function as such in humans; the DNA binding region does not bind DNA, and no promoters have been identified that target *HSFY* specifically (Shinka et al. 2004; Kichine et al. 2012). Yet, earlier reports have suggested alterations to *HSFY* expression are associated with maturation arrest of spermatogenic cells (Sato et al. 2006). Consequently, the biological functions of *HSFY* remain something of a black box in humans, let alone other species.

Looking across mammalian genomes, *HSFY* orthologues can be found from marsupial mammals to eutherian mammals, and the gene appears to be identifiable even in birds, with the inference that it has important roles in at least some species (Kinoshita et al. 2006). Mammalian *HSFY* seems present in low but variable copy number across many species, with between two and eight copies in cats (Pearks Wilkerson et al. 2008; Murphy et al. 2006), at least one retroposed active copy in mice (Kinoshita et al. 2006) and the two active copies in humans (Tessari et al. 2004) plus several pseudogene copies, one of which is found on chromosome 22. In cattle, the gene family has amplified to at least 70 copies (Hamilton et al. 2011; Yue et al. 2014). Recent work suggests that the amplification in cattle occurred after their divergence with sheep, and so is likely an independent amplification to that in pigs (Chang et al. 2013).

Gene amplifications on the sex chromosomes are particularly interesting; recent work in mice has linked the amplification of gene families on both the X and Y chromosomes to an ongoing genomic conflict affecting X gene expression and ultimately sex ratio skewing (Cocquet et al. 2012). Given the evolutionary pressures on the sex chromosomes, and the homogametic chromosome in particular, genomic conflicts are likely to be widespread across species with sex chromosome systems. Deletions on the mouse Y chromosome are known to generate reduced fertility and sperm head shape abnormalities, linked to an ongoing genomic conflict between the sex chromosomes (Ellis et al. 2011). This phenotype could be recapitulated by targeted deletion of the mouse autosomal gene *Hsf2* (Åkerfelt et al. 2008), demonstrating that these classes of transcription factors can adopt key roles in chromatin organisation. The human autosomal homologue of *HSF2* is also associated with defects in spermatogenesis (Mou et al. 2013). The classical description of the heat shock family of transcription factors is that they bind heat-shock response elements in gene promoters and activate transcription in conditions of heat or other stresses. It is clear though that the activity of heat-shock genes is not limited to stress responses, and they have important roles in development and gametogenesis (Abane and Mezger 2010).

The potential independent amplification of *HSFY* in both pigs and cattle is intriguing, as this suggests the gene is ‘prone’ to amplification on mammalian Y chromosomes, either driving or carried along with a genomic conflict. This, plus the association with fertility, makes the *HSFY* genes an important family to characterise. We show that in pigs, *HSFY* is amplified on the short arm of the chromosome, in two variant forms, with at least 100 copies combined. Both forms are expressed in testis, though only one is likely to produce a functional product. We also find both variants in other suid species, with differing copy numbers suggesting independent amplifications during suid diversification.

## Results

### Pigs carry at least two forms of *HSFY*

As the sequencing of pig Y chromosome fosmids progressed (see Skinner et al, in submission), it became clear that some sequences were present at a high copy number. BLAST searches of these sequences against the NCBI nt database suggested that one class of repetitive sequence involved the pig homologues of the *HSFY* genes. Based on the alignment of fosmid clones identified with *HSFY* copies, we designed primers to amplify fragments across both exons and the intron over all these known copies. These are shown in Table S1 as primer set 1; the schematic diagram of the *HSFY* gene and the regions amplified are also given in Figure 1. Using these primers for PCR on genomic DNA from male and female domestic pigs, we obtained distinct sequences of two lengths in the male DNA, dubbed the ‘long’ and ‘short’ forms (see Figure 2 genomic controls). The PCR products were subcloned and sequenced, and the products showed 99% sequence identity to sequence annotated within the fosmid clones. Representative examples of long and short forms can be found in the VEGA database with the accessions OTTSUSG00000005615 and OTTSUSG00000005190 respectively. We also found three instances of a third form of the sequence, longer still, supported by a faint band in the gels (not shown), exemplified by accession OTTSUSG00000002741.

**Figure 1.**
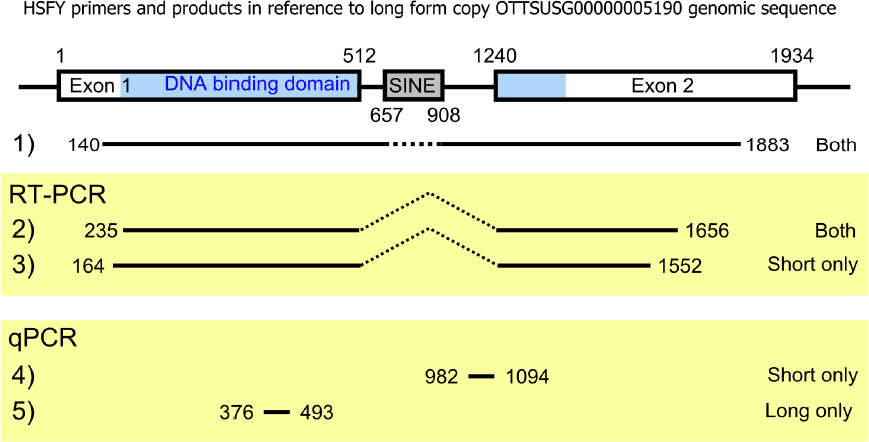
The structure of pig *HSFY*. The structure of *HSFY*, presented for the long form copy as detected in *Sus scrofa* Y-fosmid WTSI_1061-5E11 (OTTSUSG00000005190). Also shown are the regions and variants (long / short) amplified by the primer sets in Supplementary Table S1. Three of the annotated *HSFY* copies also contain a second inserted SINE within the intron.

**Figure 2.**
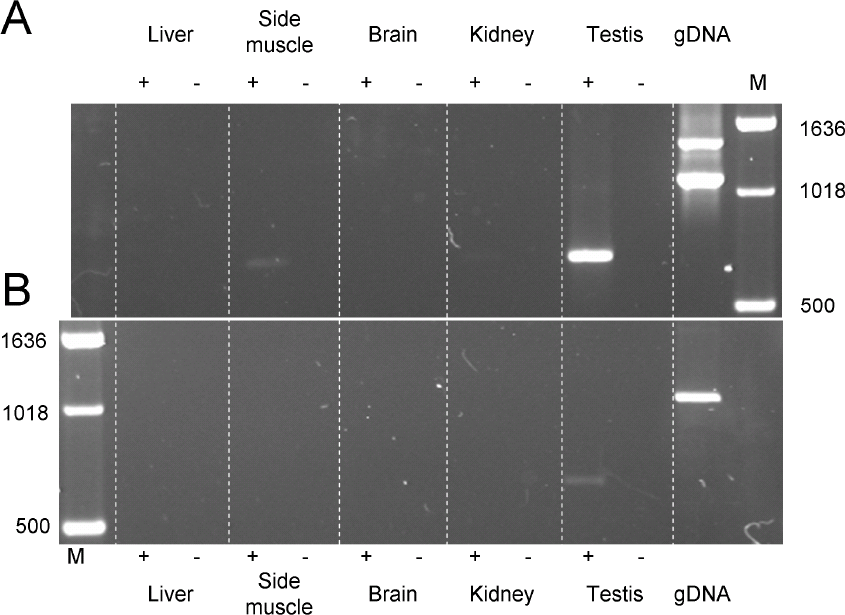
Expression status of *HSFY*. RT-PCR results on *Sus scrofa* male mRNA and gDNA using A) primer set 2 (*HSFY*exons 1 and 2 from both short and long form) and B) primer set 3 (*HSFY* exons 1 and 2 from just the short form). Tissues and DNA were taken from a Duroc male. The short form specific PCR shows expression only in testis. The pan primers show stronger expression in testis, and potentially also weak expression in side muscle. Male genomic DNA controls show long and short forms detected in A, and short form only in B.

All the genomic sequences from were aligned. The alignment showed the sequences fell primarily into two distinct clusters (*OTTSUSG*- sequences in Figure 3), with a key differentiator being an insertion of sequence within the intronic region of the longer form. Examination of the inserted sequence shows it to be a pig-lineage SINE, Pre0_SS (De Sapio et al. 2010). The third longer form contained two SINEs within the intron.

**Figure 3.**
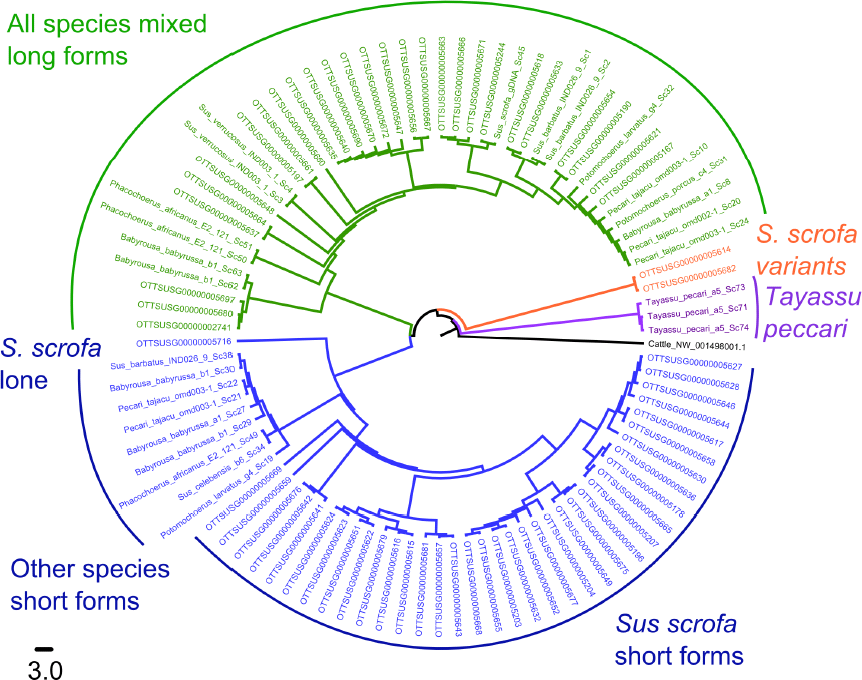
Maximum likelihood tree of *HSFY* sequences. *HSFY* copies identified through annotation of *S. scrofa* fosmids are shown by their VEGA accession number. Other suid sequences are identified by species. The corresponding region of cattle *HSFY2* was used as an outgroup. One peccary species has a distinct *HSFY* sequence; the other species all have distinct long and short forms. Long form copies show little species specific clustering amongst the fosmid sequences, whereas the short form copies show a distinct separation between *S. scrofa* fosmids and other suids. This likely reflects the potential functional nature of the short form versus the non-functional long form.

### Sus scrofa has more than 100 copies of *HSFY*

In order to estimate the number of copies of *HSFY* in the domestic pig genome, we designed qPCR primers that would amplify specifically from the ‘long’ copies or from the ‘short’ copies (Primer set 2 and 3, Table S2). The results were normalised against *SRY* copy number. This has long been assumed to be a single copy gene in the domestic pig, but we have recently found it to be present in two copies (Skinner et al, in submission). Based on an *SRY* copy number of two, and with strong caveats that this assumes ideal PCR efficiency and equal signal from all amplicons, we detected around 68 long form and 42 short form copies of *HSFY*.

Multi-copy genes are a common feature of Y chromosomes, either tandemly repeated or dispersed across the chromosome. In order to determine the physical organisation of *HSFY* on the pig Y, we performed FISH using four fosmids known to contain *HSFY* copies. These fosmids co-localised on the short arm of the Y in metaphase chromosome spreads (Figure 4A). FISH on extended DNA fibres showed that the fosmids (and the *HSFY* sequences they contain) are dispersed within this region amongst other, as-yet unidentified, sequences (Figure 4B). The total size of the *HSFY*-containing block, estimated from cytogenetic measurements, is about 5Mb.

**Figure 4.**
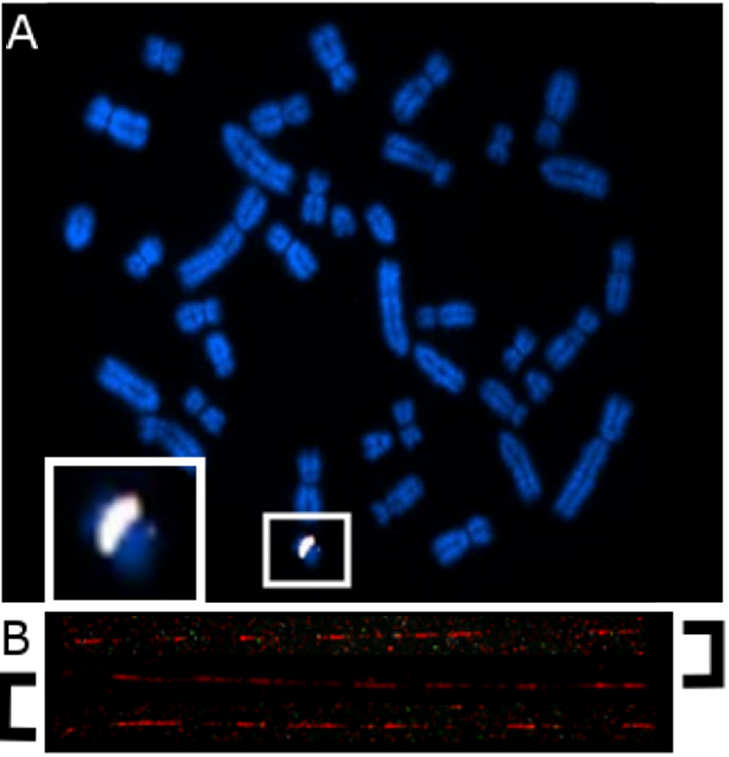
FISH using *HSFY*-containing fosmids. A) Multi colour FISH using *HSFY*-containing *Sus scrofa* WTSI_1061 Y-fosmid clones 50E19, 57F7, 69M14, 70O20 on *Sus scrofa* metaphases. Each clone is labelled with a different colour, and co-localising probes show a white signal. All the clones hybridise to the same region of the short arm of the Y chromosome (expanded in box), concordant with cytogenetic band Yp1.2. B) Fibre-FISH using fosmid clone 25019. The single 40kb clone hybridises across the ∼500kb region within the figure. Note that the hybridisation pattern is not continuous - there are other as-yet-unidentified amplified sequences within the *HSFY* region.

### Expression of *HSFY*

Multi-copy genes are of particular interest if they are expressed, especially if their expression is restricted to a certain tissue or cell type. We used reverse transcriptase PCR (RT-PCR) to characterise expression patterns of the long and short forms of *HSFY* in a range of tissues from the same animal. The primers amplified the bulk of exons 1 and 2, and the two sets designed were able to amplify either the short variants specifically, or both long and short variants together (Primer sets 2 and 3, Table S1, Figure 1). The RT-PCRs showed some expression from the short form in testis. More expression in testis was seen with the ‘both form’ primers, suggesting higher expression from the long form. The ‘both form’ primers also suggest some low levels of expression in side (loin) muscle.

At the sequence level, the long form copies all have multiple sequence changes disrupting the open reading frame, making it unlikely that a functional protein product is produced. In contrast, most of the short form copies appear to retain an open reading frame, and all but one of the identified pig ESTs matching our *HSFY* sequences cluster with the short form copies (see Supplementary Figure 1).

### *HSFY* variants in other suiforms

In order to provide information on the date of the amplification of *HSFY*, we investigated a range of related species (Table 2) with the primer sets we had available. Primer set 1 was able to amplify products from all the suids we tested, and sequencing of these products revealed both long and short forms were present in all individual animals we tested of each species, with the exception of *T. pecari*. However, current assessment of suid phylogeny (Gongora et al. 2011) has the *T. pecari* / *P. tajacu* split occurring after the divergence from the Sus lineage. Consequently, it seems more likely that these sequences represent a different evolutionary history in the *T. pecari* lineage, and not an ancestral state. This places the initial divergence between the long and short forms - i.e. the insertion of the SINE - before the diversification of modern suids (Figure 3). The peccary sequences also cluster outside the two variant sequences in *Sus scrofa*.

The qPCR primers were also used to estimate the copy number of the detectable forms of *HSFY* in each species. Again, normalisation was performed in reference to *SRY*. Although we know *Sus scrofa* has two copies of *SRY*, we do not know when this duplication occurred. Hence the error bars on the cross-species qPCR results include both possibilities (Figure 5). Conservatively considering all possible combinations of single or dual-copy *SRY*, some patterns emerge:

1. The *Sus* genus has a consistent high copy number (∼100 copies) of *HSFY*.
2. *B. babirusa* has a low copy number due to lack or amplification, or amplification of variants undetectable here, and may represent the ancestral state.
3. Warthogs and bushpigs have considerable difference between them that may be attributable to different levels of ongoing expansion of *HSFY* in each lineage.

**Figure 5.**
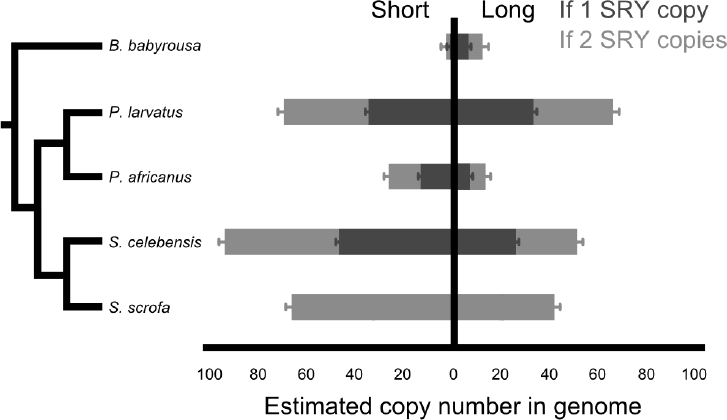
*HSFY* copy number estimates.

**Figure 6.** Estimated copy number of *HSFY* in suid species measured relative to *SRY*. Two *SRY* copies are present in S. *scrofa; SRY* copy numbers in other species are unknown, hence estimates are shown for one and two copies. In both cases, there is a clear difference between high *HSFY* copy number species and low copy number species; this can be explained by two separate amplifications - one in the *P. larvatus* lineage, and one in the *Sus* lineage. Note that estimates assume 100% PCR efficiency and equal signal from all amplicons, and cannot detect any copies with variation at the primer binding sites. Full data is given in Table 3.

### Selection within potentially coding *HSFY* copies

None of the long form copies of *HSFY* found in *Sus scrofa* or any of the other suids had coding potential. They all have frameshift and nonsense mutations occurring within the DNA-binding domain of the product rendering them likely non-functional. Given that that there is considerable gene expression from the long forms in domestic pig, it is possible that the transcripts are simply noise, or that the transcripts do not produce a protein product, but the RNAs have acquired a new function. Since there appears to be no sequence preservation in the different suid lineages (as evidenced by the ‘mixed’ clustering of the long form products), transcriptional noise seems the most likely explanation.

We therefore looked at the short form copies with a viable open reading frame, and tested for evidence of positive or purifying selection. The results of these tests are given in supplementary tables S3 and S4, and revealed that there is little evidence for positive selection amongst *HSFY* copies. There was more significant evidence for purifying selection, both within *Sus scrofa*, and between species.

## Discussion

### *HSFY* genes are amplified on pig *Y_p_*

Amplified genes and gene families are a common feature of Y chromosomes in mammals - indeed, of sex chromosomes in general. The *HSFY* genes are an example of this in pigs. We have shown here that there are two forms of *HSFY*, long and short. Both forms are present at high copy number on the Y chromosome, almost entirely located within a single cytogenetic band on the short arm. Expression analysis reveals that both forms are expressed, though evidence from EST libraries and our sequencing suggests that only the short forms are functional.

The major structural difference between the long and short forms is the presence or absence of a SINE within the intron. This SINE - *Pre0_SS* - is annotated in Repbase as being a still active pig lineage specific tRNA SINE (De Sapio et al. 2010). Given that we can find long and short forms in all the suiform species in this study, it is probable that the SINE originally inserted when there were a small number of *HSFY* copies in the ancestral genome, and subsequently both long and short copies underwent amplification. Given that we found two copies that do not cluster with long or short form, it is likely that there are other variants of *HSFY* not detected by our primer sets.

Estimates of the overall copy number of *HSFY* (Table 3, Figure 5) suggest that there are about 100 copies in the domestic pig genome, split between long and short forms, with a bias toward the short form. This number is based on comparison to *SRY*, which until recently was believed to be a single copy gene in suids (see Skinner et al). The other four species presented may also have two *SRY* copies, but the estimates we generate consider both possibilities. The other *Sus* member, *S. celebensis*, has 70 or 140 copies; again with a bias towards the short form. This suggests there has been only limited expansion of the *HSFYs* in either lineage from their common ancestor. The babirusa is an outgroup to the other species, and only a small number of copies were detected (1-2 short, 6-12 long). This suggests that either babirusa has significant copy number loss or sequence divergence, or that the *HSFY* amplifications predominantly occurred after the babirusa lineage diverged from other suids. The remaining two species tested provide tentative support to the latter scenario: the warthog *P. africanus* has a low number of copies (20-40), compared to the bushpig *P. larvatus* (70-140). Based on the phylogeny of these species, the most plausible explanation for this pattern is an amplification within the bushpig lineage. Consequently, even with the caveats of *SRY* copy number and broadness of primer coverage, there is evidence supporting two independent bursts of *HSFY* amplification within the suids.

The study here has focussed on the *HSFY* genes. However, the FISH analysis has demonstrated that the ∼5Mb *HSFY* region of the Y chromosome is not solely composed of *HSFY* copies. The full extent of the other sequences within this region is not known, due to the difficulty of assembling highly repetitive sequences reliably. Still, there are two other identified genes close to *HSFY* copies that are also amplified, thought to be pseudogenes (*RPS2* and *XKR3-like;* see also our companion Y paper for the complete context of the pig Y).

Amplified genes on the sex chromosomes have been associated with genomic conflicts in mice (e.g. Coquet et al. 2012). These genes generally act by favouring the transmission of the chromosome on which they reside, or by suppressing the transmission of their opposite gametologue (Ellis et al. 2011). The situation in pig is different to known genomic conflict models, however, in that there are no observed gene family expansions on the X chromosome that might be responding to the expansion on the Y (see Skinner et al), and we therefore consider that a similar mechanism of genomic conflict is unlikely. The X-chromosome homologue of *HSFY, HSFX*, was previously predicted (Genbank: XM_005654314.1). As with *HSFX/HSFY*comparisons in other species (e.g. Hamilton et al., 2011), there is little sequence identity between the X and Y copies. Indeed, the only alignable region is the DNA binding domain. It is clear that if there is any biological role for *HSFY*, it has been distinct from *HSFX* for the majority of mammalian Y chromosome evolution.

A further possibility is that the expansion is evolutionarily neutral - a concentration of repetitive material provided a substrate for process such as non-allelic sister chromatid exchange, causing sequence amplification, but without any selective pressure, or a biological function associated with the increase. This seems less likely; if there were no functional role for the extra *HSFY* copies, we would expect to see an accumulation of mutations within both short and long forms, abolishing the open reading frame. However, the short form copies appear to be predominantly translatable. The status of the promoters is not clear, given we have sequences for only a subset of the total *HSFY* complement: weakened or disrupted promoter activity could ‘normalise’ expression to the level of a single gene copy (and this would be consistent with the apparently lower expression levels we found from the short form (Figure 3). Further work is required to distinguish between these possibilities.

Our tests for evidence of selection for rapid amino acid change suggested no evidence for such positive selection. However, there was strong evidence for purifying selection amongst the coding copies between *Sus scrofa* and the other species, and between the *Sus scrofa* copies themselves. This again supports the idea that the copy number of these genes is functionally relevant, and that this function is maintained amongst the suid species studied here.

### Further *HSFY* variants may be present

Two *Sus scrofa* non-coding *HSFY*variants lack a SINE, but also do not cluster with the short form copies (OTTSUSG00000005614 and 5682; orange in Figure 3). These variants have nucleotide differences within the binding regions for the primer sets 1, and could not be detected in expression or copy number studies in *Sus scrofa* or any other species. It is thus possible that these are two representatives of a further diversification of the *HSFY* family; the sampled fosmids cover only a small portion of the complete ∼5Mb *HSFY*-block.

One species showed a different organisation to the others. This was the white-lipped peccary, *Tayassu pecari*. Neither the consensus short form nor the long form was identified. Instead, three similar variant species-specific forms were seen (purple sequences in Figure 4). None of these appear to have coding potential, nor is it known what the copy number of *HSFY* is in any of the peccary species. Both peccary species share a common ancestor after the divergence with the suids approximately 40 million years ago. Since *P. tajacu* has at least one each of long and short forms, it is most likely that there has been little amplification in the peccary lineage, and species-specific diversification of the *HSFY* copies in *T. pecari*. Previous comparative chromosome painting studies have suggested that the peccaries have higher rates of chromosomal rearrangement than suids (Bosma et al. 2004). Of the two peccary species in this study, the *T. pecari* karyotype appears the more derived (Adega et al. 2007, 2008), and this may contribute to the differences seen in *T. pecari*.

### A single *HSFY* pseudocopy lies outside the main block near *TSPY*

One of the copies of *HSFY* (OTTSUSG00000005716) we detected in domestic pig lies outside the *HSFY*-block, close to *TSPY* (Skinner et al, in submission). It has a premature stop codon within the DNA binding domain of the first exon, and thus cannot form a valid *HSFY* product, nor do we have evidence to suggest it is expressed. The sequence is similar to the short form, but clusters distinctly outside the other short forms (Figure 3). Its presence could be attributable to (1) an ancestral *HSFY* copy (many other species have multiple *HSFY* copies, and perhaps one of these copies gave rise to the long and short forms while the other remained unamplified; or (2) this is derived from another short form copy that relocated from the *HSFY* block during the evolution of the pig Y chromosome. We reconstructed the series of rearrangements on the Y chromosome from the ancestral mammalian Y as described in our associated X and Y sequencing paper (Skinner et al, in submission), but see no obvious opportunity for an *HSFY* copy to be relocated to the vicinity of *SRY* or *RBMY*. This does not preclude more complex undocumented rearrangements. Further cross-species cytogenetics will be able to investigate this possibility.

### Comparison with cattle suggests independent amplifications

Cattle also have a documented expansion of *HSFY* (Hamilton et al. 2011; Yue et al. 2014), also with no apparent corresponding *HSFX* expansion on the X chromosome. This opened the possibility that the amplification predated the bovine/suid divergence, and was then maintained in each lineage. Recent evidence from sheep has suggested that this is not the case, the cattle *HSFY* expansion occurring after sheep and cattle diverged about 22 million years ago (Chang et al. 2013), with variation in *HSFY* copy number between different cattle breeds (Yue et al. 2014). Accordingly, our alignments of pig *HSFY* sequences to documented cattle *HSFY* sequences show no evidence for the intronic SINE that distinguishes the long and short forms, and which must predate the initial amplification of the copies in pig. As a result, there are multiple lines of evidence pointing to independent amplifications of *HSFY* in these two lineages.

Further to this, our qPCR data provide tentative support for at least two separate amplifications of *HSFY* within the suids: once within the *P. larvatus* lineage, and again in the *Sus* lineage. However, this is subject to uncertainties of *SRY* copy number in each species and variation in qPCR primer binding sites; full confirmation of the copy numbers will require a more detailed sequencing approach to detect all variants of *HSFY* in each species.

From an evolutionary perspective, recurrent amplifications are very interesting; we do not know if the *HSFY* expansion is neutral, driven by chance and the genomic landscape within which they occur, or subject to selection for increased copy number, with an important biological role. Multiple independent amplifications of *HSFY* in different lineages suggest that there is a biological role for the gene to be discovered in pigs. In humans, the active *HSFY* genes are expressed in Sertoli cells and spermatogenic cells, potentially with a different role in each (Shinka et al. 2004); in cattle, *HSFY* is expressed in spermatogonial and spermatocyte cells (Hamilton et al. 2011). The evidence from cattle breeds shows an inverse correlation between *HSFY* copy number and testicular size, and a positive correlation with conception rate (Yue et al. 2014). It is likely that similar phenotypes will be associated with the genes in pigs. Furthermore, testicle size in pigs is correlated with the levels of the hormone androstenone in body fats (Aldal et al. 2005), which is predominantly genetically determined (Bonneau 1998). High levels of androstenone contribute to an unpleasant odour in male carcasses called boar taint, and currently lead to early castration of piglets. Consequently, further understanding the associations of *HSFY* genes with fertility and testis development will be of particular interest to the animal breeding industries.

## Conclusion

Y chromosomes are hotspots of evolutionary innovation and diversity, and it is becoming clear with increasing number of sequenced Y chromosomes that the evolutionary pressures on the sex chromosomes can drive the amplification of particular genes with dramatic functional consequences. It remains to be seen whether *HSFY* has a functional role driving its expansion, or if it has been carried as a by-product of some other process in pig and other species. Nonetheless, it appears that some genes are predisposed to amplification by their roles, locations or both.

## Methods

### Amplification of *HSFY* from genomic DNA

Primers were designed against *HSFY* sequence from pig Y fosmids. The fosmid sequences were previously deposited in Genbank as part of the pig X and Y sequencing efforts described elsewhere (Skinner et al, in submission). Primer set 1 (Supplementary Table 1) was tested on male and female gDNA from *Sus scrofa*, and on male genomic DNA from the other species in Table 2; the sampled species have been previously described (Cliffe et al. 2010), save *S. verrucosus* and *P. tajacu* DNA, which were provided by Lawrence Schook (University of Illinois). PCR followed standard protocols using Taq polymerase and buffer kit (Roche). Cycling conditions were 95°C for 3mins, 25 cycles of 95°C/ 56°C/72°C 30s for 30s/30s/60s, followed by 72°C for 10mins.

**Table 1:** Suiform species in this study. Peccaries are members of the family Tayassuidae; all other species are of the family Suidae. For a recent phylogeny of the suids, see Gongora et al, (2011). A single animal from each species was studied.

| Binomial name | Common name |
| --- | --- |
| *Sus scrofa* | Domestic pig (Duroc) |
| *Sus celebensis* | Sulawesi pig |
| *Sus verrucosus* | Java warty pig |
| *Sus barbatus* | Bornean bearded pig |
| *Potamochoerus larvatus* | Bushpig |
| *Potamochoerus porcus* | Red river hog |
| *Phacochoerus africanus* | Warthog |
| *Babyrousa babyrussa* | Buru babirusa |
| *Tayassu pecari* | White-lipped peccary |
| *Pecari tajacu* | Collared peccary |

**Table 2:** Results of qPCR on five species of long and short form copy number relative to *SRY* (primer sets 4-6 in Table 1). Since *SRY* copy number is uncertain outside *S. scrofa*, absolute values for *HSFY* are given for one and two *SRY* copies. See also Figure 6.

| Species | Copy number relative to *SRY* (+/- SEM) |  | Absolute *HSFY* copy number estimate |  |
| --- | --- | --- | --- | --- |
|  | Short | Long | Short | Long |
| *Sus scrofa* | 34 (1.09) | 21 (1.25) | 68 | 42 |
| *Sus celebensis* | 47 (1.11) | 25 (1.18) | 47 / 94 | 25 / 50 |
| *Potamochoerus larvatus* | 35 (1.15) | 33 (1.18) | 35 / 70 | 33 / 66 |
| *Phacochoerus africanus* | 13 (1.09) | 6 (1.05) | 13 / 26 | 6 / 12 |
| *Babyrousa babyrussa* | 1 (1.09) | 6 (1.09) | 1 / 2 | 6 / 12 |

### Subcloning and sequencing

PCR products were ethanol precipitated and subcloned into pGEM®-T Easy Vector System I (Promega) following the manufacturer’s instructions. Ligations were carried out at 16°C overnight. Ligated products were transformed in to XL1-Blue Competent cells (Agilent) following the manufacturer’s protocols, plated on LB-ampicillin plates supplemented with x-gal and IPTG, and incubated overnight at 37°C. White colonies were selected and used for PCR with SP6 and T7 primers to confirm product insertion. PCR products were purified following agarose gel electrophoresis and then sequenced at the sequencing facility in the Department of Genetics, University of Cambridge. Prior to the sequencing reactions, the PCR products were purified using ExoSAP-IT (USB Corporation, USA) following the manufacturer’s recommendations (samples were held at 37°C for 30 min, 80°C for 15 min, and chilled at 4°C until removed from the machine). Amplicons were sequenced using Big Dye version 3.1 (Applied Biosystems). The sequencing program consisted of 30 cycles of: 96°C/55°C/60°C for 10s/5s/4min. Products were run on an ABI 3100 capillary sequencer. Traces were edited using Chromas version 2.2 (Technelysium Pty Ltd). Sequences from different species were viewed using the MultAlin program (http://prodes.toulouse.inra.fr/MultAlin/MultAlin.html), and within the ClustalW2 program (http://www.ebi.ac.uk/Tools/clustalw2/). Repetitive content within sequences was analysed using RepeatMasker (Smit et al. 1996). Sequences have been uploaded to GenBank under accessions KP211992-KP212018. *HSFY* copies from fosmids in the domestic pig were annotated as part of the pig X and Y chromosome sequencing project (Skinner et al, in submission).

### Alignments and evolutionary analysis

Analysis of *HSFY* sequences was performed in MEGA6 (Tamura et al. 2013). The evolutionary history was inferred using the Maximum Likelihood method based on the Tamura 3-parameter model (Tamura 1992). The tree with the highest log likelihood (-13948.7067) is shown in Figure 3. Initial tree(s) for the heuristic search were obtained automatically by applying Neighbor-Join and BioNJ algorithms to a matrix of pairwise distances estimated using the Maximum Composite Likelihood (MCL) approach, and then selecting the topology with superior log likelihood value. A discrete Gamma distribution was used to model evolutionary rate differences among sites (5 categories (+*G*, parameter = 0.9667)). Branch lengths are measured in the number of substitutions per site. The analysis involved 94 nucleotide sequences. All positions with less than 95% site coverage were eliminated. That is, fewer than 5% alignment gaps, missing data, and ambiguous bases were allowed at any position. There were a total of 1433 positions in the final dataset.

ESTs corresponding to *HSFY* were identified in the NCBI EST database using BLAST (CV866737, CV873904, CX058656, CX063068, EW632312, EW633910, EW636148). Cattle *HSFY* sequence copies were identified from BLAST searches of cattle genome sequences, GenBank: NC_016145.1 (Elsik et al. 2009), using the *HSFY* sequence determined by Hamilton et al. (2012), and trimmed to match the regions of *HSFY* covered in our analysis. All sequences were aligned as above, and a phylogenetic tree was constructed using the same parameters. Newick format files for both trees are provided in Supplementary Files 1 and 2.

The 35 *HSFY* nucleotide sequences with a potentially valid open reading frame were examined separately for evidence of selection. The intronic sequences were removed, and all sequences were trimmed to the region amplified by primer set 1. The test statistics (d_S_ - d_N_) were calculated for each pairwise sequence comparison, and are given in supplementary tables 3 and 4. Analyses were conducted using the Nei-Gojobori method (Nei and Gojobori 1986).

### Expression analysis of *HSFY*

*HSFY* expression was tested only in *Sus scrofa*. Six tissues were acquired from the same boar used in the pig Y chromosome sequencing project (Skinner et al, in submission) and stored at −80C in RNALater (Qiagen). Tissues were homogenised in Trizol, nucleic acids were extracted with phenol-chloroform and DNase treated. RNA was precipitated with isopropanol and stored at 1μg/μl in ddH_2_O at −80°C. RT-PCR was carried out using a OneStep RT-PCR kit (Qiagen) on 25 ng of total RNA. PCRs used primer sets 2 and 3, given in table 1.

### qPCR

Copy number estimates were generated by quantitative PCR (qPCR). Primers (sets 4 and 5, Table 1) were designed to amplify ∼100bp fragments of the long form or the short from as wide a range of species as possible based on our sequenced products. Five species were covered: *S. scrofa, S. celebensis, P. larvatus, P. africanus* and *B. babyrussa.* Control primers (set 6, Table 1) were designed against *SRY*, presumed to be single copy, targeted to amplify from all these five species. qPCR was performed using an iCycler (BioRad) and SYPR-FAST qPCR kit (Kapa Biosystems) on male gDNA. Cycling conditions were 95°C for 3mins, followed by 40 cycles of 95°C/ 57°C/72°C 30s for 10s/20s/30s. To enable use of consistent reference genes, an annealing temperature of 57°C was used for all qPCR reactions. The fluorescent signal threshold crossing point (Ct) was normalized to the (presumed single-copy) *SRY* signal to produce a normalised ΔC_t_ and an estimate of the absolute *HSFY* copy number as 2^ΔCt^. During the course of analysis it was determined that *SRY* is dual copy in *S. scrofa*; its status in other species remains undetermined. Results are presented for both possibilities in light of this.

### Preparation of single DNA-molecule fibres by molecular combing and fibre-FISH

Single-molecule DNA fibres were prepared by molecular combing (Michalet et al. 1997) according the manufacturer’s instructions (Genomic Vision) using fibroblast cells of a Duroc boar. Briefly, the cells were embedded in a low-melt-point agarose plug (1 million cells per 90μl plug), followed by proteinase K digestion, washing in 1 × TE (10mM Tris, 1 mM EDTA, pH8.0) & beta-agarase digestion steps. The DNA fibres were mechanically stretched onto saline-coated coverslips using a Molecular Combing System (Genomic Vision). To make FISH probes, purified fosmid DNAs were first amplified using a GenomePlex® Whole genome Amplification (WGA) kit (Sigma-Aldrich) following the manufacturer’s protocols, then labelled using a WGA reamplification kit (Sigma-Aldrich) using a custom-made dNTP mix as described before (Gribble et al. 2013). For the fibre-FISH approximately 500 ng of labelled DNA from each probe and 4 μg of porcine Hybloc DNA (Applied Genetics Laboratories) were precipitated using ethanol, then resuspended in a mix (1:1) of hybridisation buffer [containing 2×SSC, 10% sarkosyl, 2M NaCl, 10% SDS and blocking aid (Invitrogen)] and deionised formamide (final concentration 50%). Coverslips coated with combed DNA fibres were dehydrated through an 70%, 90% and 100% ethanol series and aged in 100% ethanol at 65°C for 30 seconds, followed by denaturation in an alkaline denature solution (0.5M NaOH, 1.5M NaCl) for 1-3 minutes, three washes with 1×PBS (Invitrogen) and dehydration through an 70%, 90% and 100% ethanol series. The probe mix was denatured at 65°C for 10 minutes before being applied onto the coverslips and the hybridisation was carried out in a 37°C incubator overnight. The post-hybridisation washes consisted of two rounds of washes in 50% formamide/2×SSC (v/v), followed by two additional washes in 2×SSC. All post-hybridisation washes were done at 25°C, for 5 minutes each. Digoxigenin-11-dUTP (Roche) labelled probes were detected using a 1:100 dilution of monoclonal mouse anti-dig antibody (Sigma-Aldrich) and a 1:100 dilution of Texas Red-X-conjugated goat anti-mouse IgG (Molecular Probes/Invitrogen); DNP-11-dUTP (PerkinElmer) labelled probes were detected using with a 1:100 dilution of Alexa 488-conjugated rabbit anti-DNP IgG and 1:100 dilution of Alexa 488-conjugated donkey anti-rabbit IgG (Molecular Probes/Invitrogen); biotin-16-dUTP (Roche) labelled probes were detected with one layer of 1:100 dilutions of Cy3-avidin (Sigma-Aldrich). After detection, slides were mounted with SlowFade Gold® mounting solution containing 4’,6-diamidino-2-phenylindole (Molecular Probes/Invitrogen). Images were visualised on a Zeiss AxioImager D1 microscope. Digital image capture and processing were carried out using the SmartCapture software (Digital Scientific UK).

## Data access

Domestic pig Y chromosome sequences are available from Vega (http://vega.sanger.ac.uk/Sus_scrofa/Info/Index) with the accessions provided in table S2. HSFY sequences generated from other species are deposited in Genbank under accessions KP211992-KP212018.

## Acknowledgements

We gratefully acknowledge the Wellcome Trust Sanger Institute core teams for fingerprinting, mapping, archiving, library construction, sequence improvement and sequencing, Genus for providing the Duroc boar. This work was funded by BBSRC grant BB/F021372/1. The Flow Cytometry and Cytogenetics Core Facilities at the Wellcome Trust Sanger Institute and Sanger investigators are funded by the Wellcome Trust (grant number WT098051).

## Supplementary Material

**Supplementary Figure 1.**
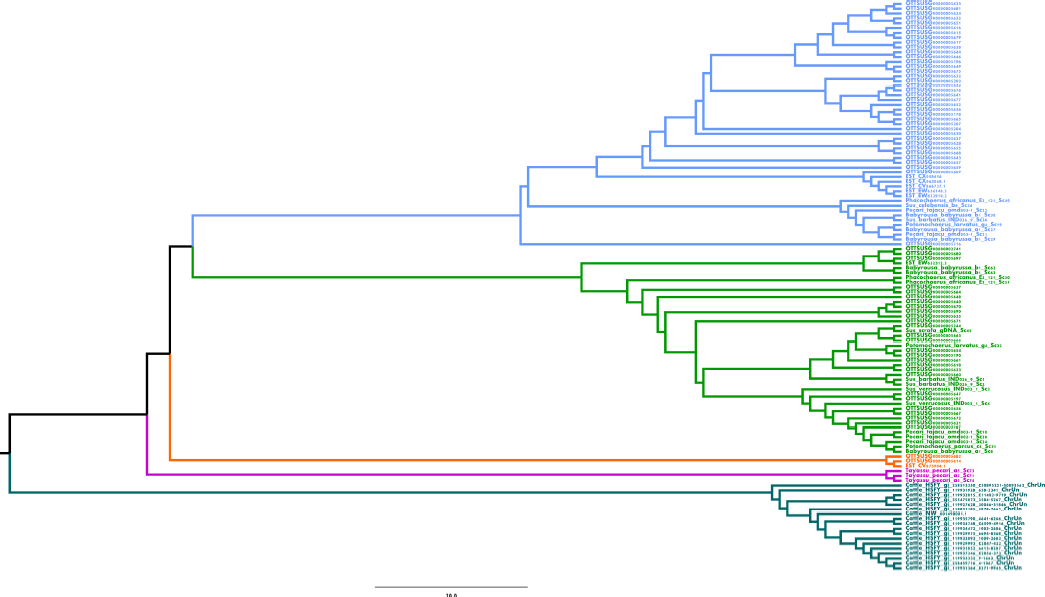
*HSFY* sequences aligned as described in Figure 3, with the inclusion of cattle *HSFY* sequences (dark blue) and pig *HSFY* ESTs (amid light blue). The clustering shows that the ESTs are almost all associated with the short form of *HSFY*, and that the cattle sequences are distinct from all suid copies, reflecting their independent amplification.

---

**Supplementary File 1 - Supplementary Tables**

**Supplementary Table S1** - Primers

**Supplementary Table S2** - Annotated *HSFY* loci on the domestic pig Y

**Supplementary Table S3** - Test for purifying selection on coding *HSFY* loci

**Supplementary Table S4** - Test for positive selection on coding *HSFY* loci

**Supplementary File 2** - Newick file of *HSFY* tree

The Newick format file containing the tree in figure 3.

**Supplementary File 3** - Newick file of complete *HSFY* tree

The Newick format file containing the tree in figure S1.

## References

Abane R, Mezger V. 2010. Roles of heat shock factors in gametogenesis and development. FEBS J 277: 4150–4172.

Adega F, Chaves R, Guedes-Pinto H. 2007. Chromosomal evolution and phylogenetic analyses in Tayassu pecari and Pecari tajacu (Tayassuidae): tales from constitutive heterochromatin. J Genet 86: 19–26.

Adega F, Chaves R, Guedes-Pinto H. 2008. Suiformes orthologous satellite DNAs as a hallmark of Pecari tajacu and Tayassu pecari (Tayassuidae) evolutionary rearrangements. Micron 39: 1281–1287.

Åkerfelt M, Henriksson E, Laiho A, Vihervaara A, Rautoma K, Kotaja N, Sistonen L. 2008. Promoter ChIP-chip analysis in mouse testis reveals Y chromosome occupancy by HSF2. Proc Natl Acad Sci 105: 11224–11229.

Aldal I, Andresen Ø, Egeli AK, Haugen J-E, Grødum A, Fjetland O, Eikaas JL H. 2005. Levels of androstenone and skatole and the occurrence of boar taint in fat from young boars. Livest Prod Sci 95: 121–129.

Bonneau M. 1998. Use of entire males for pig meat in the European Union. Meat Sci 49, Supplement 1: S257–S272.

Bosma AA, de Haan NA, Arkesteijn GJA, Yang F, Yerle M, Zijlstra C. 2004. Comparative chromosome painting between the domestic pig (Sus scrofa) and two species of peccary, the collared peccary (Tayassu tajacu) and the white-lipped peccary (T. pecari): a phylogenetic perspective. Cytogenet Genome Res 105: 115–121.

Chang T-C, Yang Y, Retzel EF, Liu W-S. 2013. Male-specific region of the bovine Y chromosome is gene rich with a high transcriptomic activity in testis development. Proc Natl Acad Sci 110: 12373–12378.

Cliffe KM, Day AE, Bagga M, Siggens K, Quilter CR, Lowden S, Finlayson HA, Palgrave CJ, Li N, Huang L, et al. 2010. Analysis of the non-recombining Y chromosome defines polymorphisms in domestic pig breeds: ancestral bases identified by comparative sequencing. Anim Genet 41: 619–629.

Cocquet J, Ellis PJI, Mahadevaiah SK, Affara NA, Vaiman D, Burgoyne PS. 2012. A Genetic Basis for a Postmeiotic X Versus Y Chromosome Intragenomic Conflict in the Mouse. PLoS Genet 8: e1002900.

De Sapio F, Jurka J, Schook LB, Archibald AL, Faulkner GJ. 2010. Non-LTR retrotransposons from pig. Repbase Rep 10:1800.

Ellis PJI, Bacon J, Affara NA. 2011. Association of Sly with sex-linked gene amplification during mouse evolution: a side effect of genomic conflict in spermatids? Hum Mol Genet 20:3010–3021.

Elsik CG, Tellam RL, Worley KC. 2009. The Genome Sequence of Taurine Cattle: A Window to Ruminant Biology and Evolution. Science 324: 522–528.

Gongora J, Cuddahee RE, Nascimento FF do, Palgrave CJ, Lowden S, Ho SYW, Simond D, Damayanti CS, White DJ, Tay WT, et al. 2011. Rethinking the evolution of extant subSaharan African suids (Suidae, Artiodactyla). Zool Scr 40: 327–335.

Groenen MAM, Archibald AL, Uenishi H, Tuggle CK, Takeuchi Y, Rothschild MF, Rogel-Gaillard C, Park C, Milan D, Megens H-J, et al. 2012. Analyses of pig genomes provide insight into porcine demography and evolution. Nature 491: 393–398.

Hamilton CK, Revay T, Domander R, Favetta LA, King WA. 2011. A Large Expansion of the HSFY Gene Family in Cattle Shows Dispersion across Yq and Testis-Specific Expression. PLoS ONE 6: e17790.

Kichine E, Rozé V, Cristofaro JD, Taulier D, Navarro A, Streichemberger E, Decarpentrie F, Metzler-Guillemain C, Lévy N, Chiaroni J, et al. 2012. HSFY genes and the P4 palindrome in the AZFb interval of the human Y chromosome are not required for spermatocyte maturation. Hum Reprod 27: 615–624.

Kinoshita K, Shinka T, Sato Y, Kurahashi H, Kowa H, Chen G, Umeno M, Toida K, Kiyokage E, Nakano T, et al. 2006. Expression analysis of a mouse orthologue of HSFY, a candidate for the azoospermic factor on the human Y chromosome. J Med Investig JMI 53: 117122.

Michalet X, Ekong R, Fougerousse F, Rousseaux S, Schurra C, Hornigold N, Slegtenhorst M. van, Wolfe J, Povey S, Beckmann JS, et al. 1997. Dynamic Molecular Combing: Stretching the Whole Human Genome for High-Resolution Studies. Science 277: 1518–1523.

Mou L, Wang Y, Li H, Huang Y, Jiang T, Huang W, Li Z, Chen J, Xie J, Liu Y, et al. 2013. A dominant-negative mutation of HSF2 associated with idiopathic azoospermia. Hum Genet 132: 159–165.

Murphy WJ, Pearks Wilkerson AJ, Raudsepp T, Agarwala R, Schaffer AA, Stanyon R, Chowdhary BP. 2006. Novel Gene Acquisition on Carnivore Y Chromosomes. PLoS Genet 2. http://www.ncbi.nlm.nih.gov/pmc/articles/PMC1420679/ (Accessed July 28, 2014).

Nei M, Gojobori T. 1986. Simple methods for estimating the numbers of synonymous and nonsynonymous nucleotide substitutions. Mol Biol Evol 3: 418–426.

Pearks Wilkerson AJ, Raudsepp T, Graves T, Albracht D, Warren W, Chowdhary BP, Skow LC, Murphy WJ. 2008. Gene discovery and comparative analysis of X-degenerate genes from the domestic cat Y chromosome. Genomics 92: 329–338.

Sato Y, Yoshida K, Shinka T, Nozawa S, Nakahori Y, Iwamoto T. 2006. Altered expression pattern of heat shock transcription factor, Y chromosome (HSFY) may be related to altered differentiation of spermatogenic cells in testes with deteriorated spermatogenesis. Fertil Steril 86: 612–618.

Shinka T, Sato Y, Chen G, Naroda T, Kinoshita K, Unemi Y, Tsuji K, Toida K, Iwamoto T, Nakahori Y. 2004. Molecular Characterization of Heat Shock-Like Factor Encoded on the Human Y Chromosome, and Implications for Male Infertility. Biol Reprod 71: 297–306.

Skinner BM, Lachani K, Sargent CA, Affara NA. 2013. Regions of XY homology in the pig X chromosome and the boundary of the pseudoautosomal region. BMC Genet 14: 3.

Tamura K. 1992. Estimation of the number of nucleotide substitutions when there are strong transition-transversion and G+C-content biases. Mol Biol Evol 9: 678–687.

Tamura K, Stecher G, Peterson D, Filipski A, Kumar S. 2013. MEGA6: Molecular Evolutionary Genetics Analysis version 6.0. Mol Biol Evol 30: 2725–2729.

Tessari A, Salata E, Ferlin A, Bartoloni L, Slongo ML, Foresta C. 2004. Characterization of HSFY, a novel AZFb gene on the Y chromosome with a possible role in human spermatogenesis. Mol Hum Reprod 10: 253–258.

Yue X-P, Dechow C, Chang T-C, DeJarnette JM, Marshall CE, Lei C-Z, Liu W-S. 2014. Copy number variations of the extensively amplified Y-linked genes, HSFY and ZNF280BY, in cattle and their association with male reproductive traits in Holstein bulls. BMC Genomics 15: 1–12.

